# Structure of KAP1 tripartite motif identifies molecular interfaces required for retroelement silencing

**DOI:** 10.1101/505677

**Authors:** Guido A. Stoll, Shun-ichiro Oda, Zheng-Shan Chong, Minmin Yu, Stephen H. McLaughlin, Yorgo Modis

## Abstract

**Abstract:** Transcription of transposable elements is tightly regulated to prevent genome damage. KRAB domain-containing zinc finger proteins (KRAB-ZFPs) and KRAB-associated protein 1 (KAP1/TRIM28) play a key role in regulating retrotransposons. KRAB-ZFPs recognize specific retrotransposon sequences and recruit KAP1, inducing the assembly of an epigenetic silencing complex, with chromatin remodeling activities that repress transcription of the targeted retrotransposon and adjacent genes. Our biophysical and structural data show that the tripartite motif (TRIM) of KAP1 forms antiparallel dimers, which further assemble into tetramers and higher-order oligomers in a concentration-dependent manner. Structure-based mutations in the B-box 1 domain prevent higher-order oligomerization without significant loss of retrotransposon silencing activity, indicating that, in contrast to other TRIM-family proteins, self-assembly is not essential for KAP1 function. The crystal structure of the KAP1 TRIM dimer identifies the KRAB domain binding site, in the coiled-coil domain near the dyad. Mutations at this site abolished KRAB binding and transcriptional silencing activity of KAP1. This work identifies the interaction interfaces in the KAP1 TRIM responsible for self-association and KRAB binding and establishes their role in retrotransposon silencing.

*Significance:* Retroviruses can integrate their DNA into the host-cell genome. Inherited retroviral DNA and other transposable elements account for over half of the human genome. Transposable elements must be tightly regulated to restrict their proliferation and prevent toxic gene expression. KAP1/TRIM28 is an essential regulator of transposable element transcription. We determined the crystal structure of the KAP1 TRIM. The structure identifies a protein-protein interaction site required for recruitment of KAP1 to transposable elements. An epigenetic gene silencing assay confirms the importance of this site for KAP1-dependent silencing. We also show that KAP1 self-assembles in solution, but this self-assembly is not required for silencing. Our work provides insights into KAP1-dependent silencing, and tools for expanding our mechanistic understanding of this process.

## Introduction

Retrovirus genomes that integrate into the genome of germline cells are inherited by future generations. These endogenous retroviruses (ERVs) can retain the ability to replicate by transcriptional amplification and expression of the viral reverse transcriptase and integrase, which convert the genome transcripts into DNA and reintegrate it into the host genome. This amplifying retrotransposition mechanism has allowed ERVs, and other retroelements such as LINEs (long interspersed nuclear elements), to accumulate, accounting for over half of the human genome (1). Approximately 100 human LINEs are still replication-competent and cause new integration events in 2-5% of the population (2).

Some ERVs and other transposable elements (TEs) have evolved to fulfill important cellular functions. TEs drive the evolution of transcriptional networks by spreading transcription factor binding sites, promoters and other regulatory elements (1, 3). TE-derived regulatory elements are particularly important in embryogenesis, when global hypomethylation promotes transcription. A significant fraction of pluripotency-associated transcription factor binding sites is located in TEs (1). TEs also serve as a reservoir of genes that can be coopted by the host. For example, TE-derived proteins catalyze V(D)J recombination (4) and syncytiotrophoblast fusion in placental development (1, 5).

Transcription of TEs must be tightly regulated, however, to prevent pathogenesis. Disruption of protein coding sequences by transposition events can cause genetic disorders such as hemophilia and cystic fibrosis (6). TE reactivation in somatic cells is associated with cancer, through disruption of tumor suppressor genes or enhanced transcription of oncogenes (6, 7). Accumulation of TE-derived nucleic acids is associated with autoimmune diseases including geographic atrophy, lupus and Sjögren’s syndrome (2, 8). A key source of retroelement repression is the family of Krüppel-associated box zinc-finger proteins (KRAB-ZFPs) and KRAB-associated protein 1 (KAP1, also known as TRIM28 or TIF1β) (9). KRAB-ZFPs, the largest family of mammalian transcription factors, recognize retroelements with a variable C-terminal array of zinc fingers (10, 11). The conserved N-terminal KRAB domain recruits KAP1 (9), which serves as a platform for the assembly of a transcriptional silencing complex of repressive chromatin-modifying enzymes including SETDB1, a histone H3K9 methyltransferase, and the nucleosome remodeling and deacetylase (NuRD) complex (12).

KAP1 is an 835-amino acid, 89-kDa protein from the tripartite motif (TRIM) family (**Fig. 1A**). Residues 57-413 contain the defining feature of the TRIM family: an RBCC motif consisting of a RING domain, two B-box-type zinc fingers and a coiled-coil domain. The RING has ubiquitin E3 ligase activity (13). This activity can be directed to tumor suppressors AMPK and p53 by the MAGE proteins, which are overexpressed in human cancers (13, 14). The resulting proteasomal degradation of AMPK and p53 has been implicated in tumorigenesis (13, 14). The central region of KAP1 contains a PxVxL motif that recruits Heterochromatin Protein 1 (HP1) and is essential for transcriptional silencing (15). The C-terminal region of KAP1 contains a PHD-bromodomain tandem (residues 624-812). The PHD recruits the SUMO E2 ligase Ubc9, to SUMOylate several lysines in the bromodomain, thereby acting as an intramolecular SUMO E3 ligase (16, 17). KAP1 SUMOylation is required for recruitment and activation of SETDB1 and recruitment of NuRD (15–17). KAP1 has also been reported to SUMOylate IRF7 (18). The KAP1 RING was necessary (but not sufficient) and the PHD dispensable for IRF7 SUMOylation (18). Similarly, the RING of PML/TRIM19 is required but not sufficient for the SUMO E3 ligase activity of PML (19, 20).

**Fig. 1.**
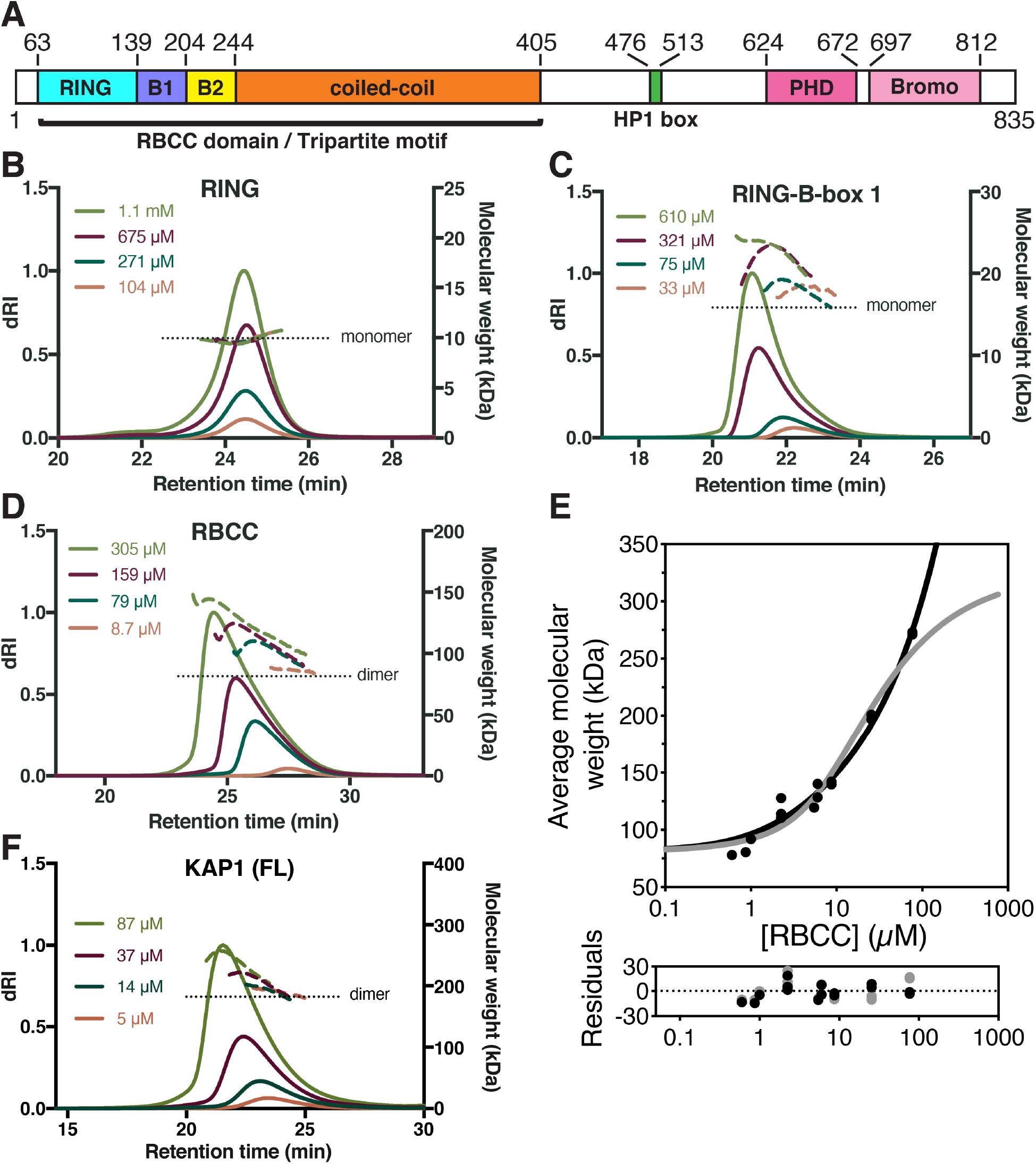
Self-assembly in solution of KAP1 RBCC and its subcomponent domains. (**A**) Domain organization of KAP1. B1, B-Box 1; B2, B-box 2. (**B-D**) SEC-MALS data for the RING, (**B**), RING-B-Box 1, (**C**), and RBCC, (**D**). (**E**) SE-AUC analysis of RBCC molecular weight as a function of protein concentration. The average molecular weight isotherm from individual fits at different concentrations was fitted to an isodesmic self-association model (black line) yielding dissociation constants of *K*_d2,4_ = 9 μM [with a 1-σ (68.3%) confidence interval of 7-11 μM] and *K*_diso_ = 19 μM [1-σ confidence interval 16-23 μM] for the dimer-tetramer and isodesmic equilibria, respectively. An alternative fit to a dimer-tetramer-octamer model (grey line) yielded dissociation constants of *K*_d2,4_ = 12.6 μM [1-σ confidence interval 7.4-23.5 μM] and *K*_d4,8_ = 6 μM [1-σ confidence interval 3-11 μM] for the dimer-tetramer and tetramer-octamer equilibria, respectively. (**F**) SEC-MALS of full-length KAP1.

The KAP1 RBCC motif has been reported to form homotrimers and bind KRAB domains with a stoichiometry of 3:1 KAP1:KRAB (21, 22). However, this is inconsistent with more recent reports that TRIM5, TRIM25 and TRIM69 form antiparallel dimers, a property predicted to be conserved across the TRIM family (23–26). Moreover, various TRIMs (TRIM5, PML/TRIM19, TRIM32) further assemble into tetramers and higher-order oligomers, including two-dimensional lattices and molecular scaffolds (as seen in PML bodies), and these higher-order assemblies are important for their biological activities (25, 27–29). RING domains, including those of TRIM5α and TRIM32, form dimers, which contribute to E3 ligase activity by priming the associated E2 ubiquitin ligase for ubiquitin transfer (28, 30, 31). The B-box 1 domain of TRIM19 (29) and B-box 2 domain of KAP1 (PDB: 2YVR) both also form dimers. Here, we use biophysical and structural approaches to show that KAP1 forms antiparallel dimers, which further assemble into tetramers and higher-order oligomers in a concentration-dependent manner. Point mutants defective in higher-order oligomerization were not significantly impaired in their retroelement silencing activity in a cell-based assay, indicating that self-assembly is not essential for the silencing function of KAP1. In contrast, mutation of conserved residues in the coiled-coil domain inhibited KRAB binding and transcriptional silencing. This work identifies the interaction interfaces in the KAP1 RBCC motif responsible for self-association and KRAB binding and establishes their role in retrotransposon silencing.

## Results

### KAP1 forms dimers that self-assemble into higher-order oligomers

Each of the domains within the RBCC motif (RING, B-box 1, B-box 2 and coiled coil) has been reported to independently form dimers or oligomers in TRIM-family proteins (23–29). The presence of two or more domains capable of oligomerization independently can lead to polymerization of TRIMs into lattices or scaffolds (25, 27, 28), but it remains unclear whether this applies to KAP1. To assess the self-assembly potential of KAP1, we purified the KAP1 RBCC motif and some of its constituent domains to determine their hydrodynamic properties. Size-exclusion chromatography coupled with multi-angle light scattering (SEC-MALS) showed that the RING domain was monomeric at concentrations ranging from 0.1 to 1.1 mM (1-10.5 g L^-1^) (**Fig. 1B**). A fragment containing the RING and B-box 1 domains was mostly monomeric at 33 μM (0.5 g L^-1^) but as the protein concentration was increased to 610 μM (9.7 g L^-1^) the average hydrodynamic radius and apparent molecular weight of the protein increased by up to 33% (**Fig. 1C**), indicating that monomers and dimers (or higher order oligomers) were in dynamic equilibrium with each other in solution. We conclude that B-box 1 drives assembly of RING-B-box 1 fragment into weakly associated dimers, with a dissociation constant in the low micromolar range.

SEC-MALS data for the whole RBCC showed unambiguously that it was dimeric at low protein concentrations (8.7 μM, 0.35 g L^-1^), and that its hydrodynamic radius and apparent molecular weight increased by up to 68% as the protein concentration was increased to 0.3 mM (12.4 g L^-1^) (**Fig. 1D**). The isolated KAP1 B-box 2 crystallized as a dimer with a surface area of 1,160 Å^2^ buried at the dimer interface (PDB: 2YVR). Together, these data suggest that the coiled-coil domain of KAP1 forms tight homodimers, like most if not all other members of the TRIM family, and that KAP1 dimers can self-assemble through further weak homotypic interactions between the B-boxes to form higher-order oligomers.

To obtain a more direct and quantitative model of KAP1 self-association we performed sedimentation equilibrium analytical ultracentrifugation (SE-AUC) on KAP1 RBCC at concentrations from 1 to 200 μM. The equilibrium sedimentation profiles were consistent with a dynamic equilibrium between dimeric and oligomeric KAP1 (**SI Appendix, Fig. S1**), in agreement with the SEC-MALS data. The average molecular mass at the lowest concentration was approximately 80 kDa (**Fig. 1E**), consistent with the formation of a dimer with a subnanomolar affinity. Further oligomerization was evident as the concentration increased. The observed increase in average molecular weight of KAP1 oligomers with increasing protein concentration at sedimentation equilibrium could be explained with two alternative models of self-association. The model with the best fit was an isodesmic self-association model in which KAP1 dimer to tetramer association is followed by unlimited consecutive additions of dimers (**Fig. 1E**). A simpler 4*R*_2_ → 2*R*_4_ → *R*_8_ model with dimers, tetramers and octamers in dynamic equilibrium produced a fit of similar quality (**Fig. 1E**). In support of the isodesmic model, the weight-average fit residuals were slightly lower at the highest protein concentrations than for the dimer-tetramer-octamer model (**Fig. 1E**). However, the improved fit of the isodesmic model could stem from the greater number of parameters versus the dimer-tetramer-octamer model. Both models yielded a dimer-tetramer dissociation constant *K*_d2,4_ and higher-order dissociation constants (*K*_d4,8_ and *K*_diso_) of the order of 10 μM. Full-length KAP1 self-assembled in a similar manner indicating that higher-order oligomerization is not an artifact of isolating the RBCC domain (**Fig. 1F**). We conclude that KAP1 forms tight dimers, which can associate into tetramers and octamers at high local concentration of KAP1. KAP1 may also form higher-order species but our SE-AUC data cannot definitively confirm or rule out the presence of KAP1 species larger than octamers.

### Crystal structure of the KAP1 RBCC tripartite motif (TRIM)

To understand the molecular basis of KAP1 self-assembly and identify its oligomerization interfaces, we determined the crystal structure of KAP1 RBCC. Although crystals of KAP1 RBCC were readily obtained, they initially diffracted X-rays poorly. Crystals suitable for structure determination were obtained by fusing bacteriophage T4 lysozyme (T4L) to the N-terminus of KAP1 RBCC and methylating primary amines in the purified protein prior to crystallization (see **Methods**). Diffraction was anisotropic, with data up to 2.63 Å resolution but with overall completeness falling below 90% at 3.9 Å resolution (**SI Appendix, Table S1**). The structure was determined by single anomalous dispersion (SAD) phasing using the anomalous scattering signal from the zinc atoms in the RING and B-boxes. The asymmetric unit contained two molecules. The atomic model was refined at 2.9 Å resolution (**SI Appendix, Fig. S2**).

The overall structure of the KAP1 RBCC dimer resembles a dumbbell (**Fig. 2**). The coiled-coil domain forms a 16 nm-long antiparallel coiled coil that contains all the dimer contacts. The ends of the coiled coil are capped by a B-box 2 domain. The RING domains (residues 63-138) are bound to one side of the coiled coil, close to but not in contact with the B-box 2 (residues 204-243) from the same subunit. Unexpectedly, there was no interpretable electron density for B-box 1 (residues 139-203), indicating that its position relative to the other domains is variable and does not obey the crystallographic symmetry. The T4L is rigidly linked to the RING domain via a continuous fused a-helix consisting of residues 158-162 from T4L (numbered 51-55 in the structure) and residues 56-62 from KAP1. The only other significant contacts between the N-terminal fusion region and the KAP1 RBCC are through the tobacco etch virus (TEV) protease cleavage site, which precedes the T4L and, atypically, is mostly ordered in the structure (**Fig. 2**). The TEV cleavage sequence is sandwiched in an extended conformation between the T4L and the coiled coil, forming multiple polar and hydrophobic contacts with both domains. Although not physiologically relevant, these contacts appear to stabilize the crystal lattice by constraining the orientation of the T4L relative to KAP1 RBCC. The T4L also forms extensive crystal packing contacts, consistent with the improved diffraction properties of the T4L-RBCC crystals versus crystals of the RBCC alone.

**Fig. 2.**
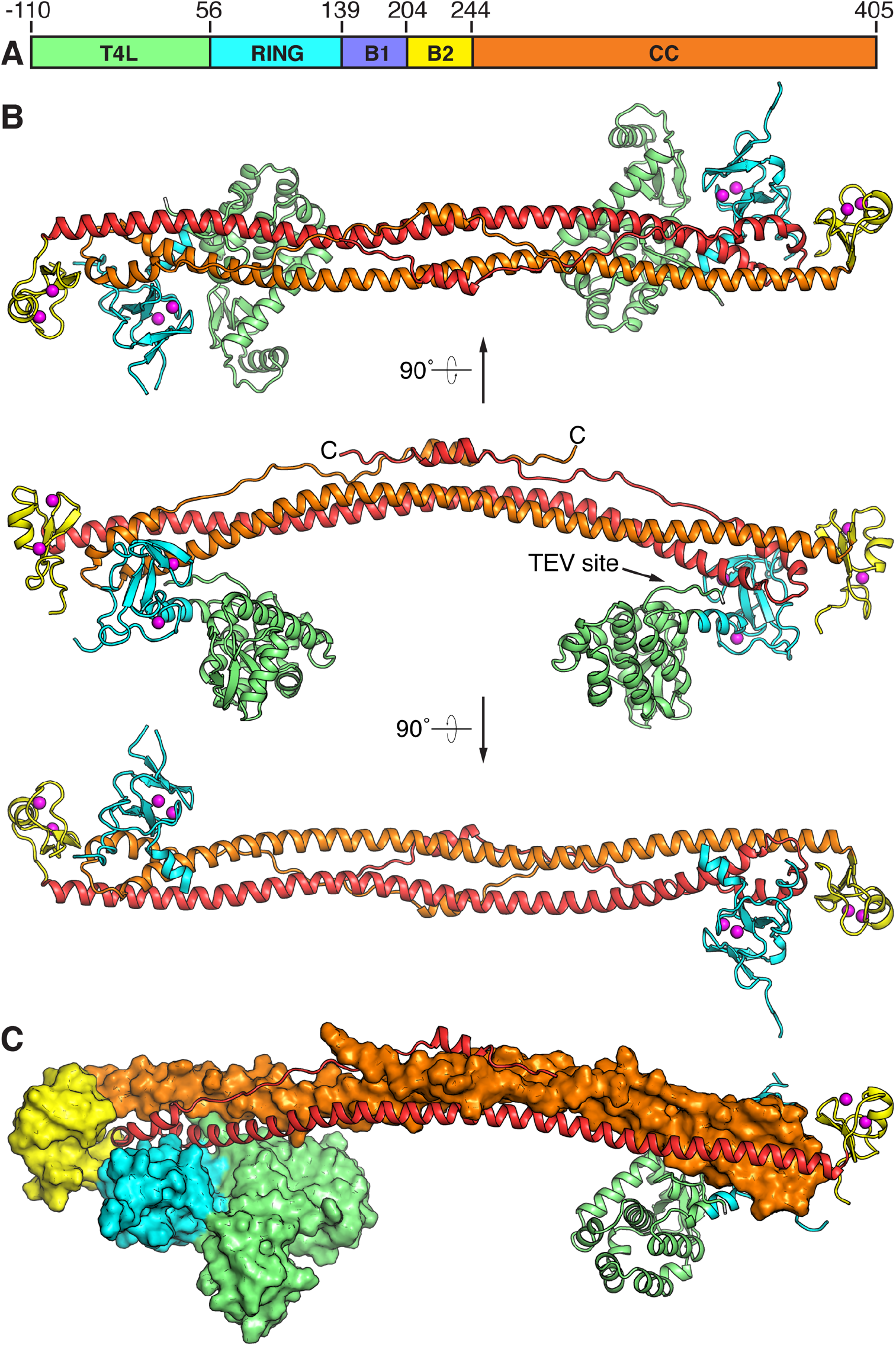
Crystal structure of KAP1 RBCC. (**A**) Domain organization of the crystallized construct. T4L, T4 lysozyme; B1, B-Box 1; B2, B-box 2; CC, coiled-coil. (**B**) Overall structure of the RBCC homodimer. Three views along or perpendicular to the dyad are shown. The components are colored as in (*A*). Zn atoms are shown as magenta spheres. (**C**) View of the RBCC dimer perpendicular to the dyad, with one subunit shown as a cartoon and the other as a surface.

The coiled-coil domain forms a helical hairpin consisting of a 15 nm-long a-helix (residues 244-348) followed by a turn and a shorter partially helical segment (residues 357-405). The first helical segment contains the majority of the dimer contacts, mostly hydrophobic leucine zipper-type coiled-coil interactions with the first helical segment from the other subunit. The second segment packs against the first to form a four-helix bundle around the twofold axis of the dimer, where the second segments from the two subunits overlap (**Fig. 2**), and a three-helix bundle at the distal ends of the dimer, where the second segments do not overlap. The central portion of the second segment has poor electron density indicating a relatively high level of conformational flexibility. The coiled-coil domain is structurally most similar to the coiled-coil domain of TRIM25 (25, 26) (Rmsd 2.6 Å), which forms a dimeric antiparallel coiled coil with the same fold and similar length and curvature. TRIM5α forms a dimeric coiled coil with the same fold and length but lower curvature (23) (Rmsd 3.9 Å), and TRIM69 forms a dimeric coiled coil with different secondary structure (24) (Rmsd 4.0 Å).

### The KAP1 RING and B-box 2 do not form dimers in the RBCC crystal structure

RING domains of E3 ubiquitin ligases recruit ubiquitin-conjugated E2 ligases to the substrate and prime ubiquitin transfer from the E2 ligase to the substrate by stabilizing the E2-ubiquitin intermediate in a closed state competent for transfer (31, 32). This stabilization has been proposed to be dependent on RING dimerization in TRIM5α, TRIM25, TRIM32 and BIRC-family E3 ligases (28, 30, 31). The most similar RING domain structure to the KAP1 RING domain is that of TRIM32 (28) (Rmsd 1.9 Å). However, in contrast to the TRIM32 RING, which dimerizes in solution via a-helices flanking the core RING domain (28), the KAP1 RING domains in the RBCC dimer are located on opposite ends of the coiled-coil domain and do not form any homotypic contacts. The KAP1 RING domains form crystal contacts with the coiled-coil domain (but not the RING domain) of a neighboring RBCC dimer. Similarly, the KAP1 B-box 2 domain also does not form homotypic contacts in the RBCC dimer. The B-box 2 domain does form crystal contacts with B-box 2 domains from two different neighboring RBCC dimers in the crystal lattice, but these homotypic contacts are distinct from those formed by the isolated B-box 2 in solution. The latter are moreover incompatible with the RBCC dimer structure as the coiled-coil domain blocks the B-box 2 surface that mediates dimerization of the isolated B-box 2.

### Higher-order assembly of KAP1 dimers is dependent on B-box 1 interactions

Our hydrodynamic data indicate that KAP1 dimers self-assemble into tetramers, octamers and higher-order oligomers through one or both of the B-boxes. To identify the sites responsible for higher-order oligomerization of KAP1, we designed structure-based mutations in the B-boxes aimed at disrupting potential dimer contacts (**Fig. 3**). In B-box 2, a cluster of residues involved in homotypic crystal contacts was mutated, yielding the variant N235A/A236D/K238A/D239A/F244A/L245A (**Fig. 3A**). Residues in the RING domain forming crystal contacts (with the coiled-coil domain) were mutated in a second variant, V114A/Q123A/F125A/K127A (**Fig. 3A**). Since B-box 1 was disordered in the RBCC structure, we mutated residues predicted to be involved in B-box 1 dimerization based on a structural model of the KAP1 B-box 1 dimer generated from the TRIM19 B-box 1 dimer structure (PDB: 2MVW) (29), yielding the variant A160D/T163A/E175R (**Fig. 3B**). The oligomerization potential of each of these variants was then assessed by SEC-MALS. Mutations in the RING and B-box 2 domains did not alter the self-assembly properties of KAP1 RBCC (**Fig. 3C and D**). The latter was unexpected since a crystal structure of the isolated KAP1 B-box 2 was dimeric (PDB: 2YVR). In contrast, the B-box 1 mutations abolished oligomerization of the RING-Box-1 fragment and almost completely inhibited higher-order oligomerization of KAP1 RBCC dimers (**Fig. 3E and F**). We conclude that the assembly of KAP1 RBCC dimers observed at high protein concentration occurs primarily through dimerization of B-box 1 domain (**Fig. 3G**).

**Fig. 3.**
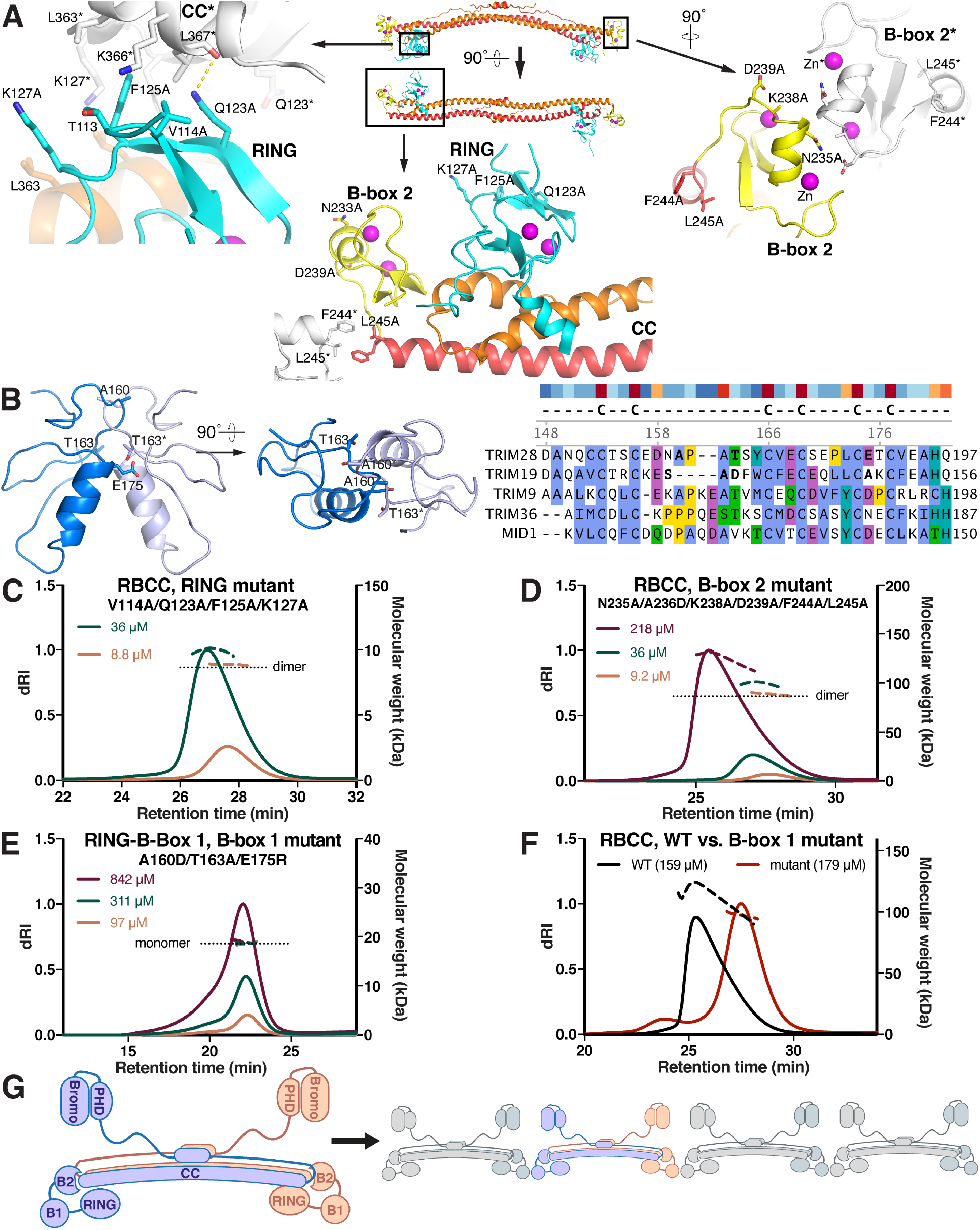
Self-assembly properties and KRAB binding activity of KAP1 RBCC mutants. (**A**) Positions of the mutations in the RING and B-box 2 domains. A reference RBCC dimer is colored as in **Fig. 2**. Adjacent RBCC dimers forming crystal packing contacts are shown in grey with their residue numbers followed by an asterisk. (**B**) Model of a KAP1/TRIM28 B-box 1 dimer based on the TRIM19 B-box 1 dimer structure (29) with selected residues forming dimer contacts shown. An alignment of B-box 1 sequences (right), the TRIM28 B-box 1 model, and the TRIM19 B-box 1 structure were used to identify mutations in KAP1 B-box 1 likely to disrupt dimer contacts. Residues known or predicted to participate in dimer contacts are shown in bold typeface in the sequence alignment. (**C-F**) SEC-MALS data for: (**C**) RBCC with RING domain mutations, (**D**) RBCC with the B-box 2 mutations, (**E**) RING-B-box 1 with B-Box 1 mutations, (**F**) RBCC (black curve, WT; red curve, B-Box 1 mutant A160D/T163A/E175R). (**G**) Model for oligomerization of KAP1 via B-Box 1.

### Self-assembly of KAP1 RBCC dimers is not required for retroelement silencing

Various TRIM proteins assemble into higher-order oligomers, two-dimensional lattices or molecular scaffolds that are important for physiological function (25, 27–29). To determine whether self-assembly of KAP1 into higher-order oligomers is required for its retroelement repression, we assayed the transcriptional silencing activities of wild type KAP1 and the oligomerization-deficient mutant. We used reporter constructs in which sequences from an SVA-D (SINE–Variable number tandem repeat–Alu, type D) retroelement (recognized by ZNF91) or a LINE-1 retroelement (recognized by ZNF93) cloned upstream of a minimal SV40 promoter strongly enhance firefly luciferase activity unless the respective KRAB-ZFP and KAP1 are both present to repress the reporter (11). The assay was adapted for use in KAP1-knockout (KO) HEK 293T cells (33), which were cotransfected with the reporter plasmid and plasmids encoding ZNF91 or ZNF93, KAP1 (WT or mutant) and *Renilla* luciferase under a constitutive promoter. Firefly luciferase luminescence from the reporter was normalized against the cotransfected *Renilla* luciferase to control for transfection efficiency. Mutations in the HP1-binding motif of KAP1 (R487E/V488E) abolished its transcriptional activity, while disruption of the PHD (C651A) resulted in moderate derepression of the reporter, consistent with previous reports (16, 34). Unexpectedly, however, the A160D/T163A/E175R B-box 1 variant had a similar repression activity as WT on the SVA and LINE-1 reporters, despite being deficient in dimer-dimer assembly (**Fig. 4**). We therefore conclude that self-assembly of KAP1 RBCC dimers into higher-order oligomers is not required for KAP1-dependent transcriptional silencing of the SVA/LINE-1 retroelements under the assay conditions.

**Fig. 4.**
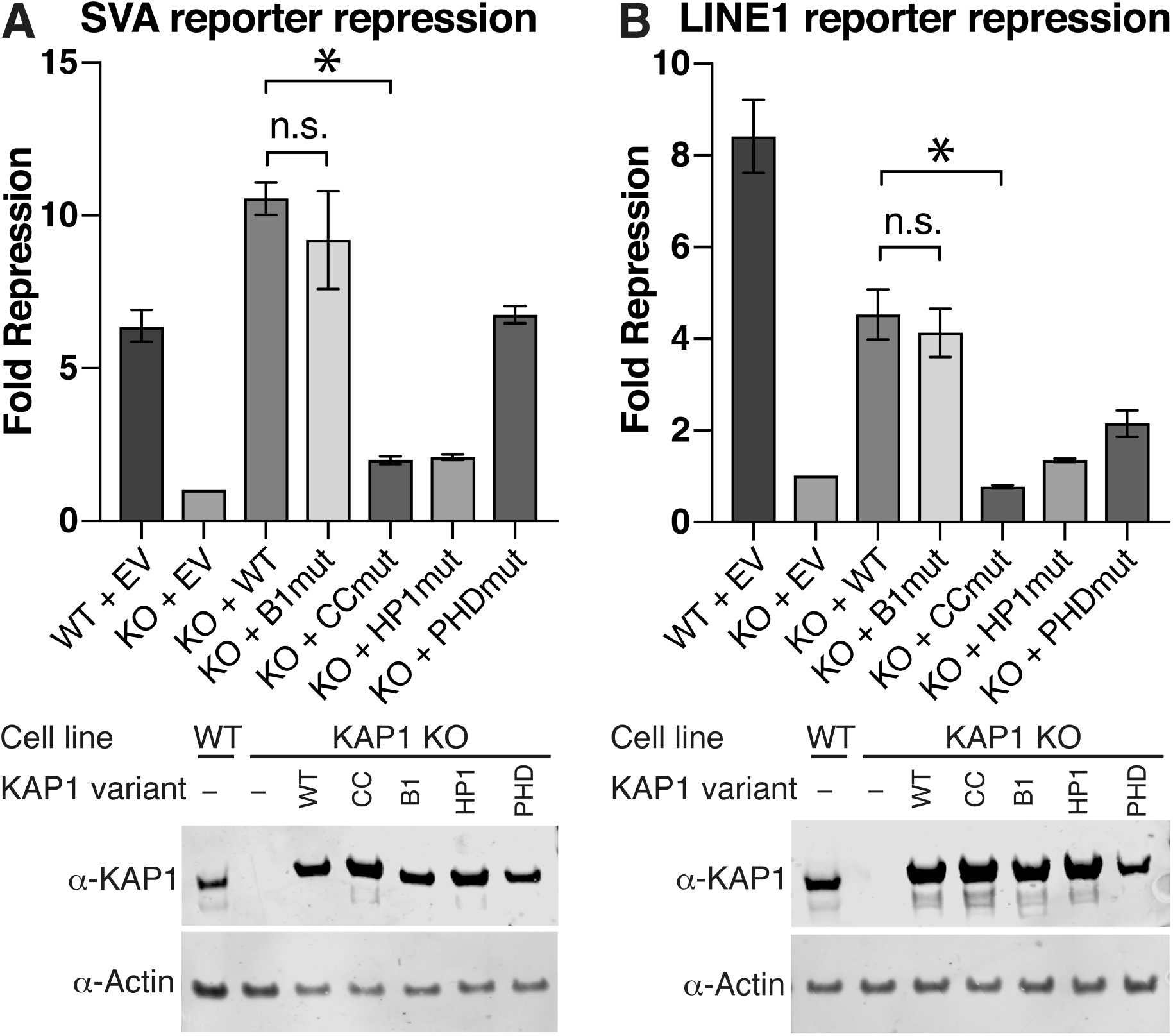
Transcriptional silencing assays with KAP1 mutants. Data are presented as fold-repression of reporter luciferase luminescence in KAP1 KO HEK293T cells transfected with a KAP1 variant. (**A**) SVA reporter repression with the oligomerization-deficient B-box 1 mutant (B1mut), the KRAB binding-deficient coiled-coil mutant (CCmut), an HP1-box mutant (HP1mut) and a PHD mutant (PHDmut). (**B**) LINE-1 reporter repression with the same set of mutants as in (**A**). Data were normalized to KAP1 KO cells transfected with an empty vector (EV). Error bars represent standard error of the mean between measurements; n = 3. Lower panels: Western blots of cell lysates from WT or KAP1 KO HEK293T cells transfected with each of the variants (WT, CC, B1, HP1, PHD) or empty vector (-).

### Conserved residues in the CC domain bind a KRAB domain and are required for silencing

The primary function of the RBCC domain of KAP1 in silencing is to bind the KRAB domains of KRAB-ZFPs and hence recruit KAP1 to its genomic targets. KRAB:KAP1 complexes were previously reported as containing one KRAB molecule and three KAP1 molecules (21, 22). However, this seemed unlikely given that KAP1 is dimeric so we decided to reexamine the composition of KRAB:KAP1 complexes. A complex of KAP1 and the KRAB domain from ZNF93, a KRAB-ZFP that binds to a LINE-1 element known to be silenced by KAP1 (11), was reconstituted by coexpressing the proteins in *E.coli.* SEC-MALS analysis showed that KAP1 and ZNF93 KRAB formed a stable complex, which retained the same ability to self-assemble into higher-order oligomers as KAP1 alone (**SI Appendix, Fig. S3A**). The average molecular mass derived from SEC-MALS for the KAP1-KRAB complex of 226 kDa was inconsistent with a 1:3 KRAB:KAP1 stoichiometry and suggested instead the stoichiometry of the complex was 1:2 KRAB:KAP1 (236 kDa theoretical molecular weight; **Fig. 5A**).

**Fig. 5.**
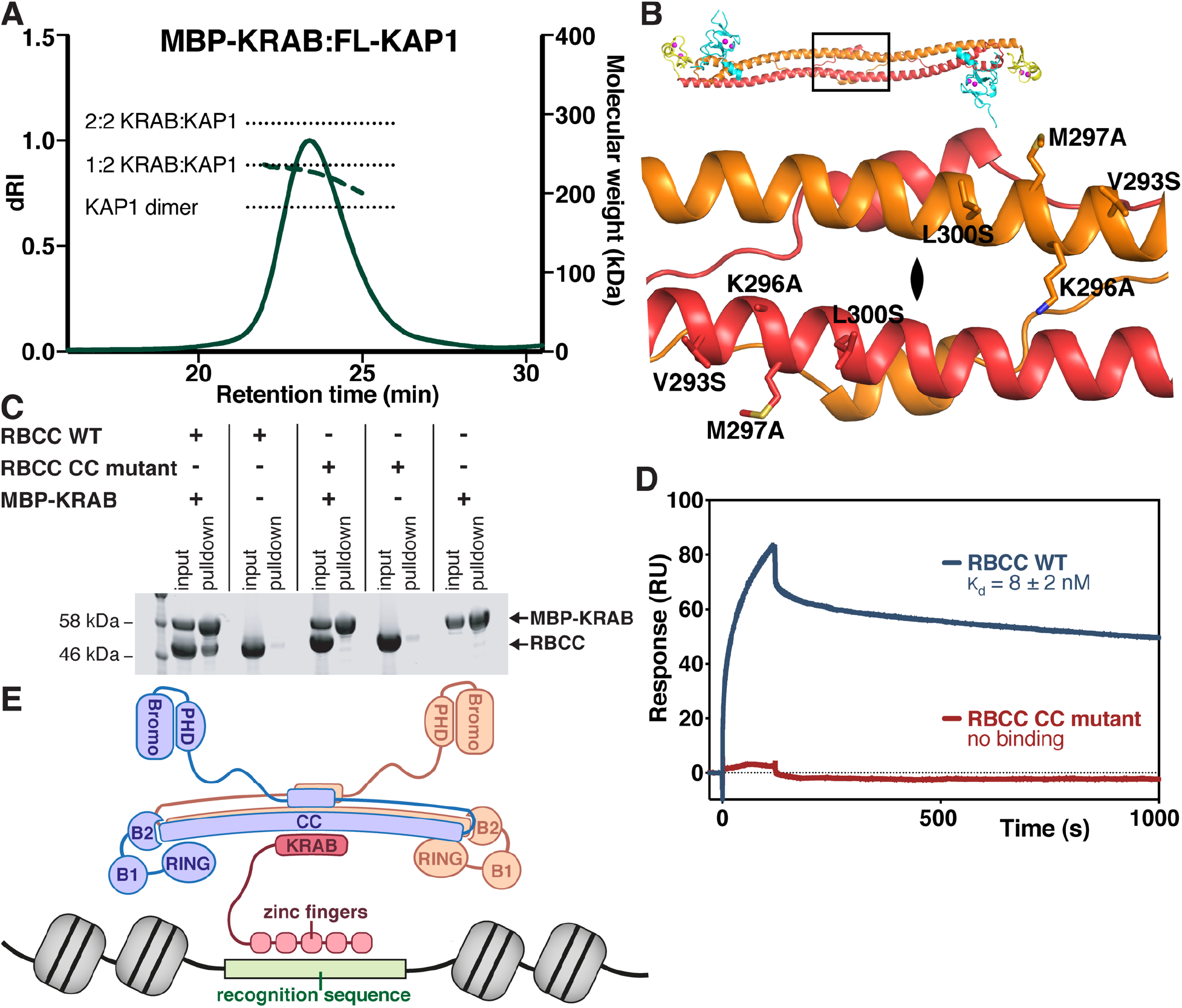
Formation of a 2:1 KAP1:KRAB complex and identification of KRAB binding residues in the KAP1 coiled-coil domain required for silencing. (**A**) SEC-MALS of full-length KAP1 bound to ZNF93 MBP-KRAB. The expected molecular weights of a KAP1 dimer, and for 2:1 and 2:2 KAP1:KRAB complexes are indicated with dashed lines. The total protein concentration of each analyte was 1.2 g l^-1^. (**B**) Closeup of the cluster of solvent-exposed hydrophobic residues near the dyad. The variant V293S/K296A/M297A/L300S (CC mutant) was generated to test for KRAB binding. (**C**) Pulldown KAP1-KRAB binding assay. KAP1 RBCC was incubated with Twin-StrepII-MBP-ZNF93 KRAB and the mixture loaded on Strep-Tactin Sepharose. Bound proteins were detected by SDS-PAGE/Coomassie. (**D**) SPR KAP1-KRAB binding assay. MBP-KRAB was immobilized on the chip. WT or CC mutant KAP1 RBCC were flowed over the chip. Binding kinetics of WT RBCC: *k*_on_ = 3.6 ± 0.96 x 10^4^ M^-1^ s^-1^; *k*_off_ = 2.7 ± 0.1 x 10^-4^ s^-1^. (**E**) Model for binding of KAP1 dimers to KRAB-ZFPs.

Having established that each RBCC dimer binds a single KRAB domain, we reasoned that the interaction interface must be located on the dyad, in the central region of the KAP1 coiled-coil domain, as every other location would result in two equivalent binding sites (and a 2:2 stoichiometry). Intriguingly, examination of our KAP1 RBCC crystal structure revealed a cluster of solvent-exposed hydrophobic residues near the twofold axis (V293, M297, L300; **Fig. 5B**). Moreover, these amino acids are conserved in KAP1 but not present in other TRIMs (**SI Appendix, Fig. S4**). To determine whether this region of the coiled-coil domain mediates KRAB binding, we designed the variant V293S/K296A/M297A/L300S (**Fig. 5B**). KAP1 RBCC domain bearing these mutations failed to bind to MBP-KRAB in a pulldown assay (**Fig. 5C**). Other properties of KAP1 such as dimerization and higher-order oligomerization were unaffected (**SI Appendix, Fig. S3B**), indicating that the mutations did not interfere with the overall fold of the RBCC domain. Notably, the thermal stability of this variant as assessed by differential scanning fluorimetry (DSF) was significantly higher than that of WT RBCC domain, further supporting a functional role of these residues (**SI Appendix, Fig. S3C**). A second variant with mutations on the dyad on opposite face of the coiled-coil domain, F391A/L395S/W398A, was mostly insoluble. The lack of binding of the V293S/K296A/M297A/L300S variant to MBP-KRAB was confirmed with surface plasmon resonance (SPR) measurements. While WT RBCC domain bound MBP-tagged KRAB domain with high affinity (8 ± 2 nM K_d_), no binding was observed with the coiled-coil mutant (**Fig. 5D** and **SI Appendix, Fig. S5**).

The effect of the coiled-coil mutations on the transcriptional silencing activity of KAP1 was measured using the reporter assay described above. Consistent with its inability to bind KRAB the V293S/K296A/M297A/L300S variant lost all repression activity, while being expressed at similar concentrations as WT KAP1 (**Fig. 4**). These data indicate that KRABs bind KAP1 on the twofold axis of the RBCC dimer on a surface that includes residues V293/K296/M297/L300 and that this binding surface is required for KAP1 repression activity.

## Discussion

Initial biophysical studies on the KAP1 RBCC domain suggested that it formed trimers, both in isolation and in complex with a KRAB domain (21, 22). More recent work on other TRIMs, in contrast, demonstrated that the coiled-coil domains of various TRIMs mediate formation of antiparallel dimers and suggests that this property is conserved across the entire TRIM family (23–26). Our SEC-MALS and SE-AUC data establish that KAP1 is dimeric rather than trimeric, and that dimers self-assemble into tetramers, octamers and possibly higher-order species at high local concentration of KAP1. Notably, the cooperativity of self-assembly implies that octamers are slightly more stable than tetramers, which could in principle promote the formation of large oligomeric assemblies of KAP1 in the nucleus. This concentration-dependent self-association of KAP1 is primarily encoded by B-box 1 and is relatively weak with a dissociation constant in the low micromolar range. The lack of a discrete endpoint oligomeric state (e.g. tetramers) led us to hypothesize that the RBCC may be able to self-associate into polymeric chains or molecular scaffold (**Fig. 3G**) as observed for other TRIMs (25, 27–29). We therefore investigated the possibility that KAP1 self-assembly may contribute to its transcriptional silencing function. Conceivably, formation of large oligomeric assemblies of KAP1 at its genomic target loci might amplify silencing activity by increasing the number of recruited repressive chromatin-modifying molecules such as SETDB1. However, KAP1 mutants deficient in self-assembly repressed transcription from SVA and LINE-1 reporter retrotransposons with an efficiency similar to WT KAP1, indicating that at least in this setting higher-order oligomerization is not absolutely required for KAP1 function.

The KAP1 RING has ubiquitin E3 ligase activity (13). E3 ligase activity is generally thought to be dependent on RING dimerization, in the context of RBCC dimerization, in TRIM5α, TRIM25, TRIM32 and BIRC-family E3 ligases (28, 30, 31). Unexpectedly, the crystal structure of KAP1 RBCC is incompatible with RING dimerization as the RING domains are positioned on opposite ends of the RBCC dimer (**Fig. 2B**). Furthermore, the KAP1 RINGs do not form significant crystal contacts and do not dimerize or self-associate in solution. The absence of homotypic contacts between the RINGs (or B-box 2 domains) in the RBCC dimer suggests that KAP1 may be amongst the minority of E3 ligases that can promote ubiquitin transfer from E2 to substrate without forming RING dimers, perhaps using structural elements from outside the core RING domain as seen for example in CBL-B (32). Alternatively, the primary function of the RING may be to contribute to SUMO E3 ligase activity, consistent with reports that the RINGs of KAP1 (18) and PML (19, 20) are required for SUMOylation of specific substrates.

The interaction between KAP1 and KRAB-ZFPs is critical for recruitment of KAP1 to retroelements, but the stoichiometry of KAP1-KRAB complexes has remained unclear and the location of the KRAB binding surface on KAP1 unknown. Our SEC-MALS data shows unambiguously that the KAP1 RBCC dimer can only bind a single KRAB domain. Moreover, we demonstrate that KRAB binding occurs on the dyad of the RBCC dimer on a surface that includes residues V293/K296/M297/L300, and that this interface is required for KAP1 repression activity. It was previously proposed that RING domain, B-box 2 and coiled-coil domain of KAP1 all contribute to KRAB binding (21, 22). However, these results were inferred from mutations in residues essential for maintaining the structural integrity of the KAP1 RBCC dimer that would likely cause misfolding of the protein. Based on the distance of the B-box 2 from the KRAB-binding residues in the coiled-coil domain and the small size of KRAB domains, direct involvement of B-box 2 in KRAB binding can be ruled out (**Fig. 5E**). Similarly, a contribution of the RING, which is also distant from the dyad, appears highly unlikely. We note that since a single KRAB domain binds to the RBCC dimer on (or near) the dyad, the contacts formed by the KRAB domain with each of the KAP1 protomers in the dimer are necessarily different, lending an inherent asymmetry to the KRAB:KAP1 interaction. Hence, loss of twofold symmetry in the KRAB:KAP1 complex is a consequence of KRAB binding to the dyad.

We have mapped the specific molecular features within the KAP1 RBCC domain that are responsible for KAP1 self-assembly and KRAB binding. We show that binding of a single KRAB domain to the dimeric antiparallel coiled-coil domain of KAP1 is the first step in KAP1-depedendent epigenetic silencing of retrotransposon transcription. Further studies, informed by the work reported here, are needed to complete our understanding of how KAP1 recognizes KRAB-ZFPs, how it activates chromatin-modifying enzymes in subsequent steps of transcriptional silencing, and how it promotes ubiquitination of specific substrate proteins without RING dimerization.

## Methods

### Expression vectors

Synthetic genes encoding KAP1 RING (residues 50-146), RING-B-box 1 (RB1; residues 50-200), RBCC (residues 50-413) and full-length KAP1 (residues 1-835; UniProt: Q13263) codon-optimized for *Escherichia coli (E. coli)* were cloned into the first multiple cloning site (MCS) of the pETDuet plasmid (Novagen), with N-terminal hexahistidine purification (His_6_) tag followed by a TEV protease cleavage site (ENLYFQG). The T4L-RBCC fusion construct was constructed by inserting a synthetic gene encoding the RBCC motif (residues 56-413) of human KAP1 codon-optimized for *E.coli* into the first MCS of pETDuet. The gene was preceded by sequences encoding: a His_6_ tag; a TEV protease cleavage site; and bacteriophage T4 lysozyme (T4L) with the N-terminal methionine deleted and the last three residues replaced by a single alanine residue. A synthetic gene encoding the KRAB domain (residues 1-71) from ZNF93 (UniProt: P35789) codon-optimized for *E.coli* was expressed from the pET20 plasmid (Novagen) with N-terminal Twin-StrepII and maltose binding protein (MBP) affinity tags (the MBP tag was also required for protein solubility). For coexpression with KAP1, the same ZNF93 KRAB construct was subcloned into MCS1 of pCDFDuet.

KAP1 RBCC and the KRAB domain of ZNF93 were coexpressed from the pETDuet plasmid. A synthetic gene encoding residues 50-413 of KAP1 codon-optimized for *E.coli* was inserted into MCS1 adding an N-terminal His6 tag followed by a TEV protease cleavage site. A synthetic gene encoding residues 1-71 of ZNF93 codon-optimized for *E.coli* was cloned into MCS2 adding an N-terminal glutathione S-transferase (GST) followed by a human rhinovirus (HRV) 3C protease cleavage site.

For the transcriptional silencing assay, expression constructs containing WT or mutant full-length human KAP1 preceded by a triple FLAG tag and linker sequence MDYKDHDGDYKDHDIDYKDDDDKGSGG were assembled in pLEXm (35) with the NEBuilder HiFi DNA Assembly Cloning Kit (New England BioLabs). The following plasmids were used: firefly luciferase reporter plasmids pGL4cp-VNTR-OCT4Enh-E2 and pGL4cp-L1PA4-OCT4Enh-E2; pCAG_ZNF91_HA encoding ZNF91; pCAG_ZNF93 encoding ZNF93; and pRTTK_Renilla encoding *Renilla* luciferase under a constitutive promoter (33).

### Protein expression and purification

*E. coli* BL21 (DE3) cells (New England BioLabs) were transformed with the respective expression construct and starter cultures were grown overnight at 30°C in 2×TY medium. Starter cultures were used to inoculate 2×TY medium and cells were incubated at 37°C and 220 rpm to an optical density (OD_600_) of 0.4-0.5. For the expression of KAP1 constructs, cultures were then supplemented with 50 μM ZnSO_4_ and the temperature of the incubator was set to 18°C. In the case of KRAB domain constructs, the incubator temperature was lowered to 16°C. Protein expression was induced at OD_600_ = 0.8 with 0.2 mM isopropyl-β-D-thiogalactopyranoside (IPTG). All subsequent steps were performed at 4°C. After 16 h cells were harvested by centrifugation, and stored at −80°C.

To purify the RBCC domain of KAP1 cells were resuspended in lysis buffer containing 50 mM Tris pH 8, 0.3 M NaCl, 20 mM imidazole, 0.5 mM TCEP, 1:10,000 (v/v) benzonase solution (Sigma), 1×cOmplete EDTA-free protease inhibitors (Roche). The cells were lysed by sonication immediately after addition of 1 mM phenylmethane sulfonyl fluoride (PMSF). The lysate was clarified by centrifugation (30 min, 40,000×g). The supernatant was applied to a 5-ml HisTrap HP nickel-affinity column (GE Healthcare) preequilibrated in wash buffer (50 mM Tris pH 8, 0.3 M NaCl, 20 mM imidazole, 0.5 mM TCEP). The column was washed with 30 column volumes (CV) of wash buffer before elution with elution buffer (50 mM Tris pH 8, 0.3 M NaCl, 0.25 M imidazole, 0.5 mM TCEP). Subsequently, the buffer was exchanged to 50 mM Tris pH 8, 0.3 M NaCl, 0.5 mM TCEP and the His_6_ tag was removed by incubating the protein overnight at 4°C with 1:50 (w/w) TEV protease. Following a second nickel-affinity chromatography step to remove uncleaved protein and protease, the sample was further purified by size-exclusion chromatography using a HiLoad (16/600) Superdex 200 pg column (GE Healthcare) preequilibrated in 20 mM HEPES pH 8, 0.5 M NaCl, 0.5 mM TCEP.

RING and RB1 constructs were purified as described above, except that a Superdex 75 (10/300) column (GE Healthcare) equilibrated in 20 mM HEPES pH 8, 0.2 M NaCl, 0.5 mM TCEP was used for the final size-exclusion chromatography step. T4L-RBCC fusion protein was purified as the RBCC domain, except that the His_6_ tag was not removed. Full-length KAP1 was purified as T4L-RBCC, except that a Superose 6 increase (10/300) column (GE Healthcare) preequilibrated in 20 mM HEPES pH 8, 0.2 M NaCl, 0.5 mM TCEP was used for the final size-exclusion chromatography step.

To purify MBP-tagged ZNF93 KRAB domain, bacteria pellets were resuspended in lysis buffer (50 mM Tris pH 8, 0.15 M NaCl, 0.5 mM TCEP, 1:10,000 (v/v) benzonase solution (Sigma), 1×cOmplete EDTA-free protease inhibitors (Roche)) and lysed by sonication. The lysate was clarified by centrifugation (30 min, 40,000×g). The supernatant was applied to a 5 ml StrepTrap column (GE Healthcare) preequilibrated in wash buffer (50 mM Tris pH 8, 0.15 M NaCl, 0.5 mM TCEP). The column was washed with 30 CV of wash buffer, before the protein was eluted with wash buffer supplemented with 2.5 mM D-desthiobiotin and further purified by size-exclusion chromatography using a HiLoad (16/600) Superdex 200 pg column (GE Healthcare) preequilibrated in 20 mM HEPES pH 8, 0.5 M NaCl, 0.5 mM TCEP. KAP1:MBP-KRAB complex was purified as the isolated KRAB domain, except that lysis and wash buffer contained 0.2 M NaCl and a Superose 6 increase (10/300) column was used for the final size-exclusion chromatography step.

### Size-exclusion chromatography and multiangle light scattering (SEC-MALS)

100 μl of protein sample was subjected to SEC at 293 K on a Superdex 200 (10/300) column (GE Healthcare) preequilibrated in 20 mM HEPES pH 8, 0.5 M NaCl, 0.5 mM TCEP (for KAP1 RBCC), a Superose 6 (10/300) column in 20 mM HEPES pH 8, 0.2 M NaCl, 0.5 mM TCEP (for full-length KAP1), a Superose 6 (10/300) column in 20 mM HEPES pH 8, 0.5 M NaCl, 0.5 mM TCEP (for KAP1:MBP-KRAB complex) or a Superdex 75 (10/300) column in 20 mM HEPES pH 8, 0.2 M NaCl, 0.5 mM TCEP (for RING and RB1 constructs) with a flow rate of 0.5 ml min-1. The SEC system was coupled to multi-angle light scattering (MALS) and quasi-elastic light scattering (QELS) modules (DAWN-8+, Wyatt Technology). Protein in the eluate was also detected with a differential refractometer (Optilab T-rEX, Wyatt Technology) and a UV detector at 280 nm (1260 UV, Agilent Technology). Molar masses of peaks in the elution profile were calculated from the light scattering and protein concentration, quantified using the differential refractive index of the peak assuming a dn/dc of 0.186, using ASTRA6 (Wyatt Technology).

### Sedimentation-equilibrium analytical ultracentrifugation (SE-AUC)

KAP1 RBCC samples at 0.2 mM (8 g l^-1^) and 12 μM (0.5 g l^-1^) were diluted in a 1:3 series in 20 mM HEPES pH 8, 0.5 M NaCl, 0.5 mM TCEP. 110 μL samples were loaded in 12 mm 6-sector cells and centrifuged at 5, 8.5 and 15 krpm at 20°C in an An50Ti rotor in an Optima XL-I analytical ultracentrifuge (Beckmann). At each speed, comparison of several scans was used to judge whether equilibrium had been reached. The data were analyzed in SEDPHAT 13b (36). Equilibrium sedimentation distributions were fit to obtain average masses. An SE-AUC average mass isotherm compiled from fits to the data was analyzed in SEDPHAT using isodesmic and dimer-tetramer-octamer oligomerization models. The partial-specific volumes (v-bar), solvent density and viscosity were calculated with Sednterp (www.rasmb.org/sednterp).

### X-ray crystallography

Prior to crystallization, free amines in T4L-RBCC were methylated by incubating 15 ml of protein solution (~1 g L^-1^) in 20 mM HEPES pH 8, 0.5 M NaCl with 300 μl of 1 M dimethylamine borane complex (ABC; Sigma-Aldrich) and 600 μl of 1 M formaldehyde for 2 h at 4°C. An additional 300 μl of 1 M ABC and 600 μl of formaldehyde were then added. After further 2 h at 4°C, 150 μl of ABC was added and the sample was incubated overnight at 4°C. The reaction was then quenched with 1.875 ml of 1 M Tris pH 8. The sample was supplemented with 2 mM DTT and purified with a HiLoad (16/600) Superdex 200 pg column preequilibrated in 20 mM HEPES pH 8, 0.5 M NaCl, 0.5 mM TCEP (37). Crystals were grown at 18°C by sitting drop vapor diffusion. Methylated T4L-RBCC at 4.5 g L^-1^ (72 μM) was mixed with an equal volume of reservoir solution optimized from the Index screen (Hampton Research): 15% (w/v) PEG 3350, 75 mM MgCl2, 0.1 M HEPES pH 7.5. Plate-shaped crystals appeared after 2 days and were frozen in liquid nitrogen with 33% ethylene glycol as a cryoprotectant. X-ray diffraction data were collected at 100 K at Diamond Light Source (beamline I03) and processed with autoPROC (38) and STARANISO (Global Phasing, Ltd). The X-ray energy was tuned to 9,672 eV, corresponding to the zinc L-III edge, for data collection. Phases were determined with the single anomalous dispersion (SAD) method in PHENIX (39) using zinc as the anomalously scattering heavy atom. Comparison of electron density maps calculated with different high-resolution cutoffs and B-factor sharpening factors (**SI Appendix, Fig. S2**) indicated that a 2.9-Å cutoff and −33 Å^2^ sharpening factor produced the best map. The atomic model was built with COOT (40) and iteratively refined with REFMAC (41) and PHENIX at 2.9 Å resolution. The atomic models for T4 lysozyme (PDB: 1LYD, 2LZM) and KAP1 B-box 2 (PDB: 2YVR) were docked into the phased electron density and the rest of the atomic model was built *de novo* using available structures of other TRIMs as guides (PDB: 4LTB (25), 4TN3 (23), 5FEY (28), 5NT1 (26)). See **SI Appendix** for data collection and refinement statistics (**Table S1**), and sample electron density (**Fig. S2**). Structure figures were generated with PyMOL (Schrodinger, LLC). Methylation of T4L-RBCC at 38 sites was confirmed by mass spectrometry (**SI Appendix, Fig. S6**), but none of the dimethyl-amine groups were visible in the map.

### Atomic model of KAP1 B-box 1

An atomic model of the KAP1 B-box 1 domain was generated from the TRIM19 B-box 1 structure (PDB: 2MVW) (29) with Phyre2 (www.sbg.bio.ic.ac.uk/phyre2). The KAP1 B-box 1 was then superimposed onto each protomer of the TRIM19 B-box 1 dimer to generate a model of the KAP1 B-box 1 dimer.

### Pulldown assay

8 nmol of KAP1 RBCC was incubated with 2 nmol of Twin-StrepII-MBP-KRAB for 45 min on ice. StrepII-tagged bait protein was then captured with 100 μl of Strep-Tactin Sepharose (IBA) for 1 h at 4°C. After four washes with 1 ml of buffer (20 mM HEPES pH 8, 0.5 M NaCl, 0.5 mM TCEP), the beads were boiled in 100 μl of 2×SDS-PAGE loading buffer and bound proteins analyzed by SDS-PAGE/Coomassie.

### Surface plasmon resonance (SPR)

SPR was performed using a Biacore T200 with dextran-coated CM5 chips (GE Healthcare). Reference control and analyte CM5 chips were equilibrated in 20 mM HEPES pH 8.0, 0.5 M NaCl, 0.5 mM TCEP at 20°C. MBP-KRAB was immobilized onto the chips until a response unit value of approximately 600 was reached. SPR runs were performed with analytes injected for 120 s followed by a 900 s dissociation in 1:2 dilution series with initial concentrations of 34 μM for WT KAP1 and 35 μM for mutant KAP1. The sensor surface was regenerated after each injection cycle with 20 mM NaOH for 30 s with a 120-s post-regeneration stabilization period. Data were fitted using a biphasic kinetic model with KaleidaGraph (Synergy Software) and PRISM 8 (GraphPad) to determine *k*_on_, *k*_off_ and *K*_d_.

### Transcriptional silencing assay

KAP1 silencing activity was measured with a reporter assays in which a SINE-VNTR-Alu (SVA) type D or LINE-1 sequence upstream of a minimal SV40 promoter enhances firefly luciferase activity unless KAP1 and the cognate KRAB-ZFP (ZNF91 and ZNF93, respectively) are present to repress the reporter (11). The assay was adapted for use in HEK293T cells as described (33). KAP1 KO 293T cells in 24-well plates were cotransfected with 20 ng firefly luciferase reporter plasmid, 0.2 μg plasmid encoding ZNF91 or ZNF93, 0.2 μg pLEXm plasmid encoding WT or mutant KAP1 and 0.4 ng plasmid encoding *Renilla* luciferase using 1.5 μl of FuGENE 6 (Promega). Luciferase activity was measured 48 h post-transfection in cell lysates with the Dual Luciferase assay kit (Promega) and a Pherastar FS platereader (BMG Labtech). Replicates were performed on separate days. Firefly luciferase values were normalized to *Renilla* luciferase values to control for transfection efficiency.

### Western blotting

4 x 10^5^ HEK293T cells were lysed in 100 μl Passive Lysis Buffer (Promega). 10 μl of cell lysates were boiled with 2.5 μl 4x gel loading buffer for 5 min at 98°C and run on a NuPAGE 4-12% Bis-Tris polyacrylamide gel for 45 min at 200 V. Gels were blotted using an iBlot Gel Transfer Device and Transfer Stacks (ThermoFisher). Blots were blocked with 5% milk in PBS for 1 h at room temperature and incubated with primary antibody overnight at 4°C. Rabbit anti-KAP1 antibody (Abcam ab10484) was diluted 1:10,000 whilst rabbit anti-actin antibody (Abcam ab219733) was diluted 1:2,000. Blots were incubated with fluorescent secondary antibodies for 30 min at room temperature. Secondary anti-rabbit antibodies were diluted 1:10,000. Blots were imaged using an Odyessy CLx gel scanner (LI-COR Biosciences).

### Statistics

For the KAP1 complementation assays, data are representative of five independent experiments and statistical significance was assessed using an unpaired t-test (assuming Gaussian distributions, without Welch’s correction) with PRISM 8. Repression activity data are represented as the mean ± s.e.m. Statistical significance was assigned as follows: n.s., P > 0.05; *, P < 0.05; **, P < 0.01, n = 3.

### Data availability

The structure factors and atomic coordinates were deposited in the Protein Data Bank (PDB: 6QAJ). The original experimental X-ray diffraction images were deposited in the SBGrid Data Bank (Dataset 637). Other data are available from the corresponding author upon request.

## Acknowledgments

KAP1 KO cells and plasmids for the transcriptional silencing assay were kind gifts from Helen Rowe (University College London). The plasmids were provided with permission from David Haussler (Univ. of California Santa Cruz). We thank Christopher Douse and other members of the Modis lab for insightful discussions. Crystallographic data were collected on beamline I03 at Diamond Light Source (DLS). Access to DLS (proposal MX15916) was supported by the Wellcome Trust, MRC and BBSRC. This work was supported by Wellcome Trust through Senior Research Fellowship 101908/Z/13/Z to YM and PhD Studentship 205833/Z/16/Z to GS.

