## Supplementary material for "Structure of KAP1 tripartite motif identifies molecular interfaces required for retroelement silencing": SI Appendix

**This PDF file includes:**

Supplementary text
Figs. S1 to S6
Table S1

### Supplementary Information Text

### Supplementary Methods

**Expression and purification of the KAP1 RBCC:KRAB complex.** KAP1 RBCC and the KRAB domain of ZNF93 were coexpressed from the pETDuet plasmid. A synthetic gene encoding residues 50-413 of KAP1 codon-optimized for *E.coli* was inserted into MCS1 adding an N-terminal $His_6$ tag followed by a TEV protease cleavage site. A synthetic gene encoding residues 1-71 of ZNF93 codon-optimized for *E.coli* was cloned into MCS2 adding an N-terminal glutathione S-transferase (GST) followed by a human rhinovirus (HRV) 3C protease cleavage site. The complex was expressed in *E. coli* BL21 (DE3) cells and cell lysates were prepared as described for the full-length KAP1:MBP-KRAB complex. The clarified lysate was applied to a 5-ml GSTrap 4B glutathione Sepharose column (GE Healthcare) preequilibrated in wash buffer (50 mM Tris pH 8, 0.15 M NaCl, 0.5 mM TCEP). The column was washed with 30 column volumes (CV) of wash buffer before elution with elution buffer (50 mM Tris pH 8, 0.15 M NaCl, 25 mM reduced glutathione, 0.5 mM TCEP). Subsequently, the buffer was exchanged to 50 mM Tris pH 8, 0.15 M NaCl, 0.5 mM TCEP and the GST tag was removed by incubating the protein overnight at 4˚C with 1:100 (w/w) HRV 3C protease. Following a second glutathione-affinity chromatography step to remove uncleaved protein and protease, the sample was further purified by size-exclusion chromatography using a HiLoad (16/600) Superdex 200 pg column (GE Healthcare) preequilibrated in 20 mM HEPES pH 8, 0.2 M NaCl, 0.5 mM TCEP.

**Differential scanning fluorimetry (DSF).** 10 µl samples containing WT or mutant KAP1 at 25-50 µM (1-2 g $L^{-1}$) in 20 mM HEPES pH 8.0, 0.5 M NaCl, 0.5 mM TCEP were loaded into glass capillaries (Nanotemper) by capillary action. Intrinsic protein fluorescence at 330 nm and 350 nm was monitored between 15 and 95°C in a Prometheus NT.48 instrument (Nanotemper), and the $T_m$ values calculated within the accompanying software by taking the turning point of the first derivative of the $F_{350}$:$F_{330}$ ratio as a function of temperature.

### Supplementary Figures and Figure Legends

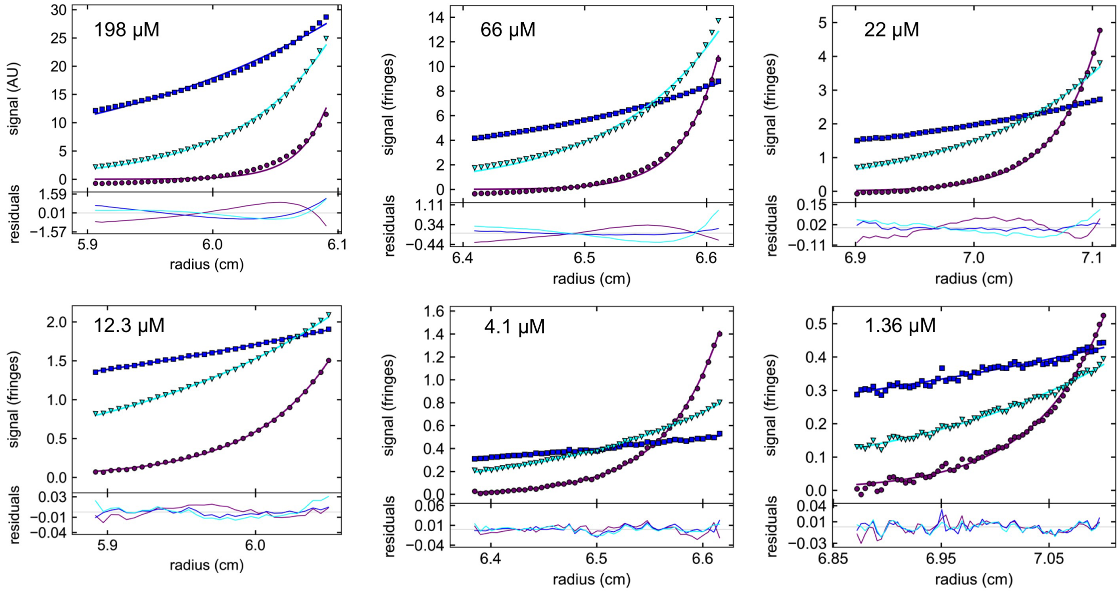


**Fig. S1.** Sedimentation equilibrium analytical centrifugation (SE-AUC) of KAP1 RBCC. The equilibrium distributions of KAP1 RBCC, at different concentrations in each inset, during centrifugation at 5,000 rpm (blue squares), 8,500 rpm (cyan triangles) and 15,000 rpm (magenta circles) were fitted to a single species model (solid lines) to obtain average molecular weights.

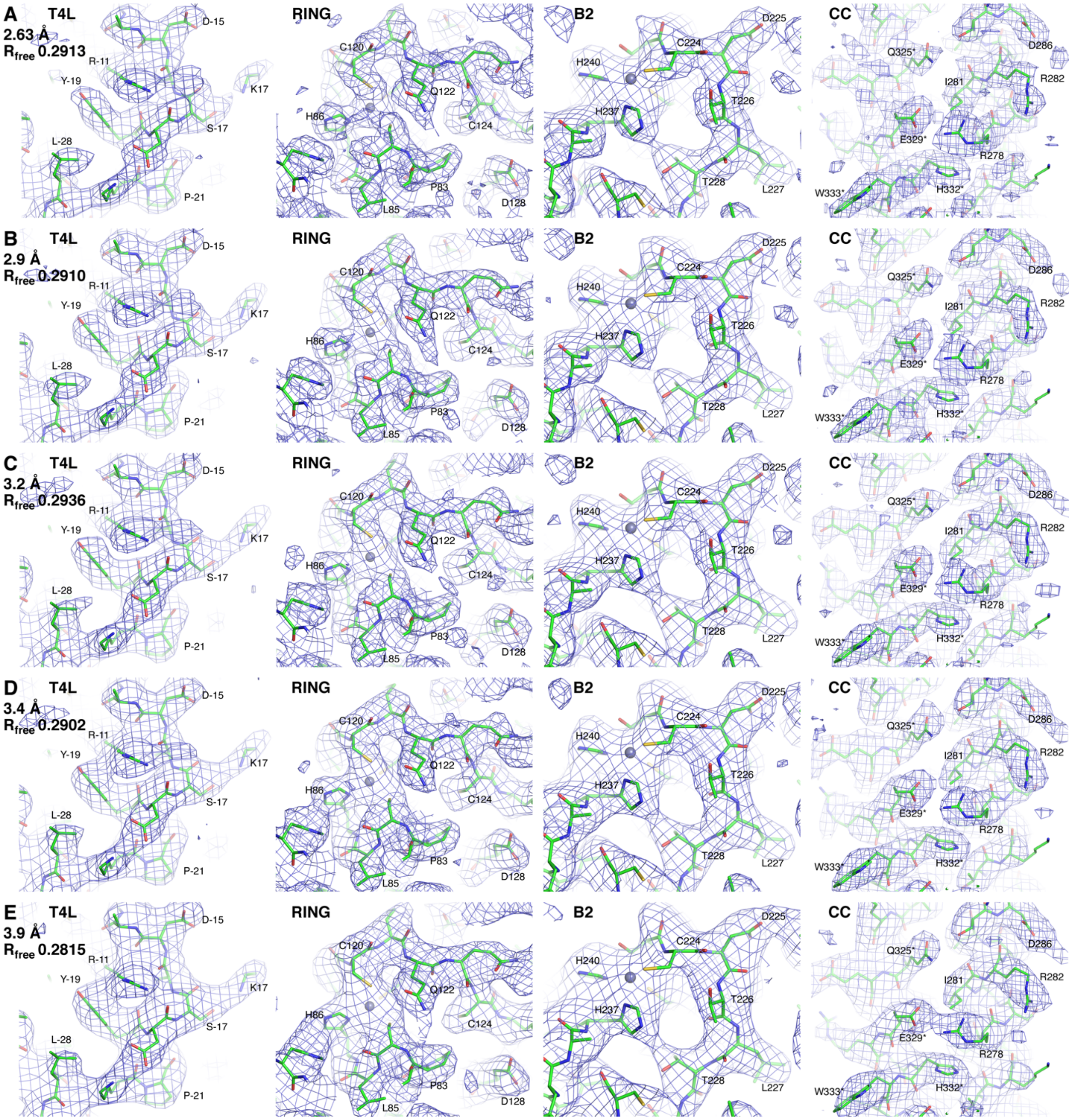


**Fig. S2.** Representative samples of electron density from the KAP1 RBCC crystal structure. Electron density maps were calculated after one cycle of refinement with Phenix v1.15 at 2.63 Å resolution, (**A**). 2.9 Å resolution, (**B**), 3.2 Å resolution, (**C**), 3.4 Å resolution, (**D**), and 3.9 Å resolution, (**E**). Electron density samples from the T4 lysozyme (T4L), RING, B-box 2 (B2) and coiled-coil (CC) domains are shown. A contour level of 1.0 σ (rmsd = 1) was used for all panels. Map sharpening factors were applied with COOT v0.8.9.2 to optimize the appearance of the maps as follows: 2.63-Å map, -43 $Å^2$; 2.9-Å map, -33 $Å^2$; 3.2-Å map, -25 $Å^2$; 3.4-Å map, -19 $Å^2$; 3.9-Å map, -7.6 $Å^2$. The 2.9-Å map was used for model building.

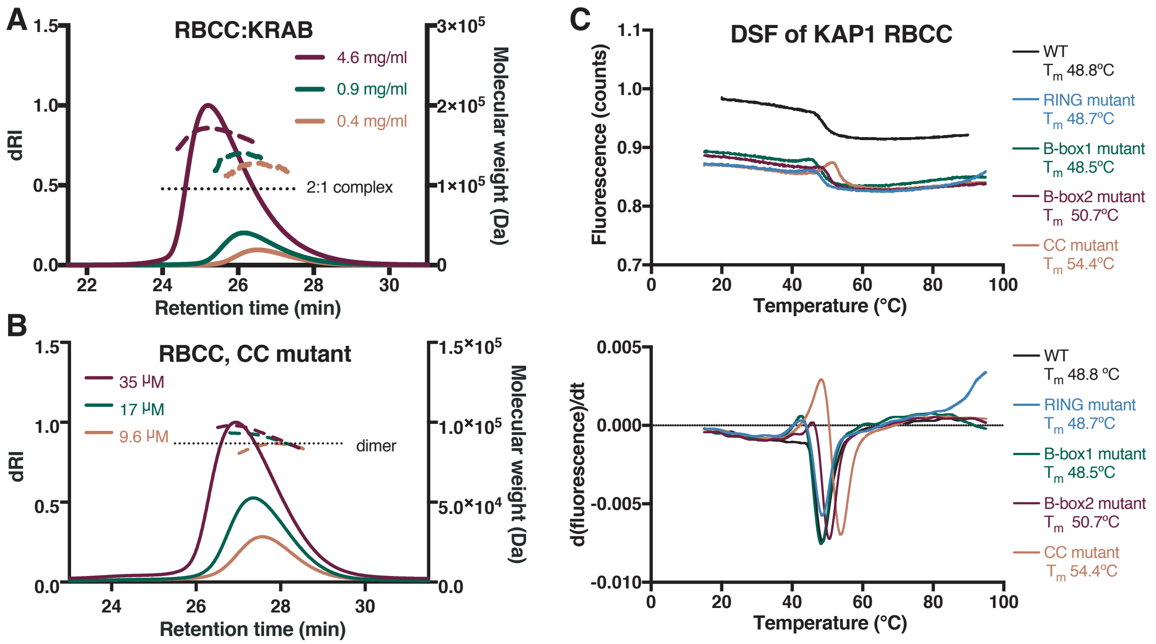


**Fig. S3.** Hydrodynamic properties and thermal stability of KAP1 RBCC variants. (**A**) SEC-MALS of RBCC-KRAB complexes at various concentrations. (**B**) SEC-MALS of the KAP1 RBCC variant with the coiled-coil domain mutations (V293S/K296A/M297A/L300S) at various concentrations. (**C**) Differential scanning fluorimetry (DSF) of the KAP1 RBCC (WT and variants). Intrinsic protein fluorescence at 330 nm and 350 nm was monitored between 15 and 95°C. Melting temperatures ($T_m$) were calculated from the turning point of the first derivative of the $F_{350}$:$F_{330}$ ratio as a function of temperature.

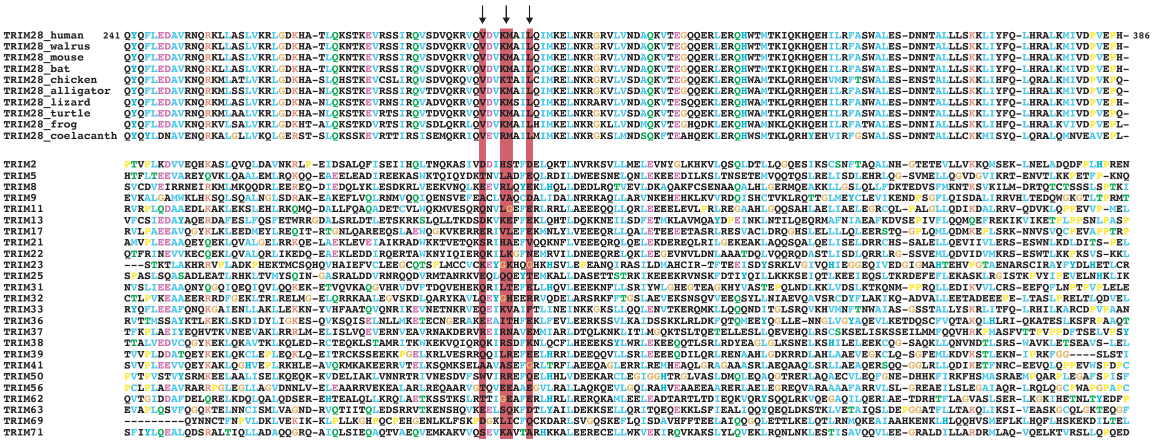

**Fig. S4.** Multiple sequence alignment of TRIM coiled-coil domains. The coiled-coil domains of KAP1/TRIM28 from various vertebrate species (upper sequence cluster), and of other human TRIMs (lower sequence cluster) are shown. Conservation of residues involved in KRAB binding is highlighted with red columns and arrows.

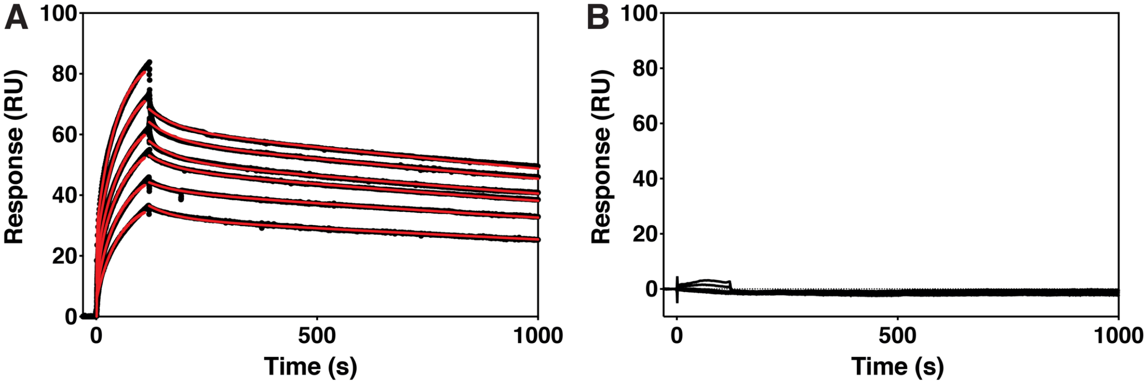


$k_{off1} = 2.7 \pm 0.1 \times 10^{-4}\ s^{-1}$
$k_{on1} = 3.6 \pm 0.96 \times 10^{4}\ M^{-1}\ s^{-1}$
$K_{d1} = 8 \pm 2$ nM

$k_{off2} = 1.6 \pm 0.6 \times 10^{-2}\ s^{-1}$
$k_{on2} = 138 \pm 60\ M^{-1}\ s^{-1}$
$K_{d2} = 118 \pm 39\ \mu M$

**Fig. S5.** Surface plasmon resonance (SPR) analysis of KAP1 RBCC. Analysis of the interactions of wild-type RBCC, (**A**), and of the coiled-coil mutant (V293S/K296A/M297A/L300S), (**B**), with MBP-KRAB immobilized on the sensor chip. Sensograms are shown for 1:2 dilution series starting from 34 µM for wild-type RBCC and 35 µM for mutant RBCC. The fits for the association and dissociation kinetics for wild-type KAP1 are shown in red with the corresponding rate constants and derived dissociation constants.

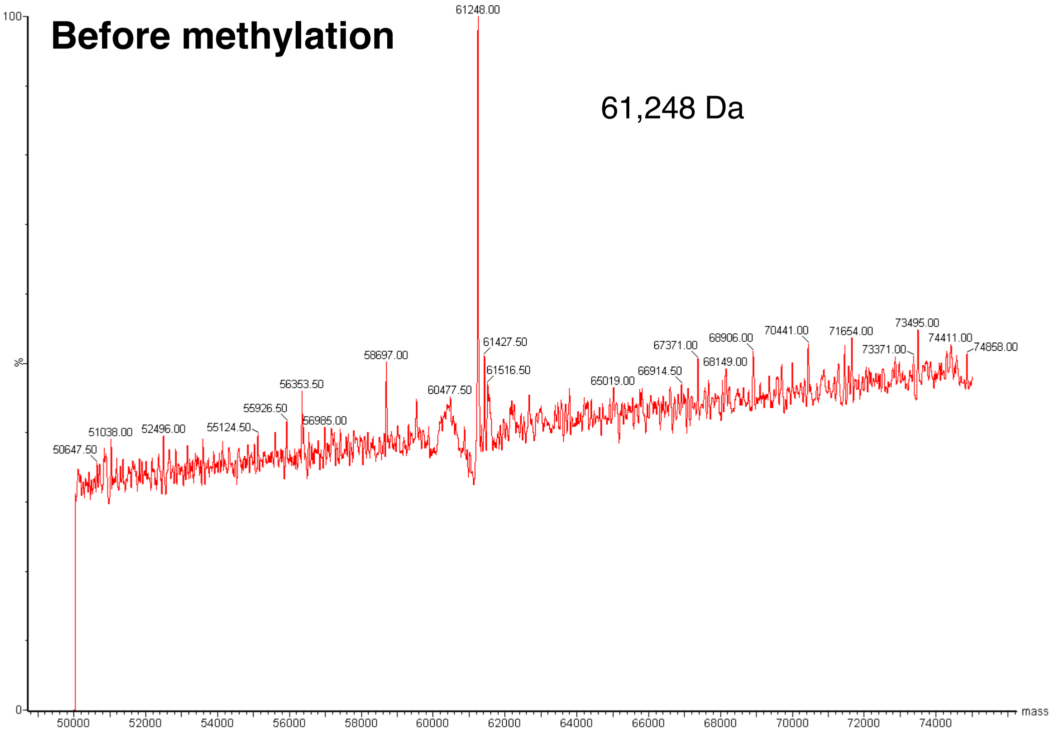


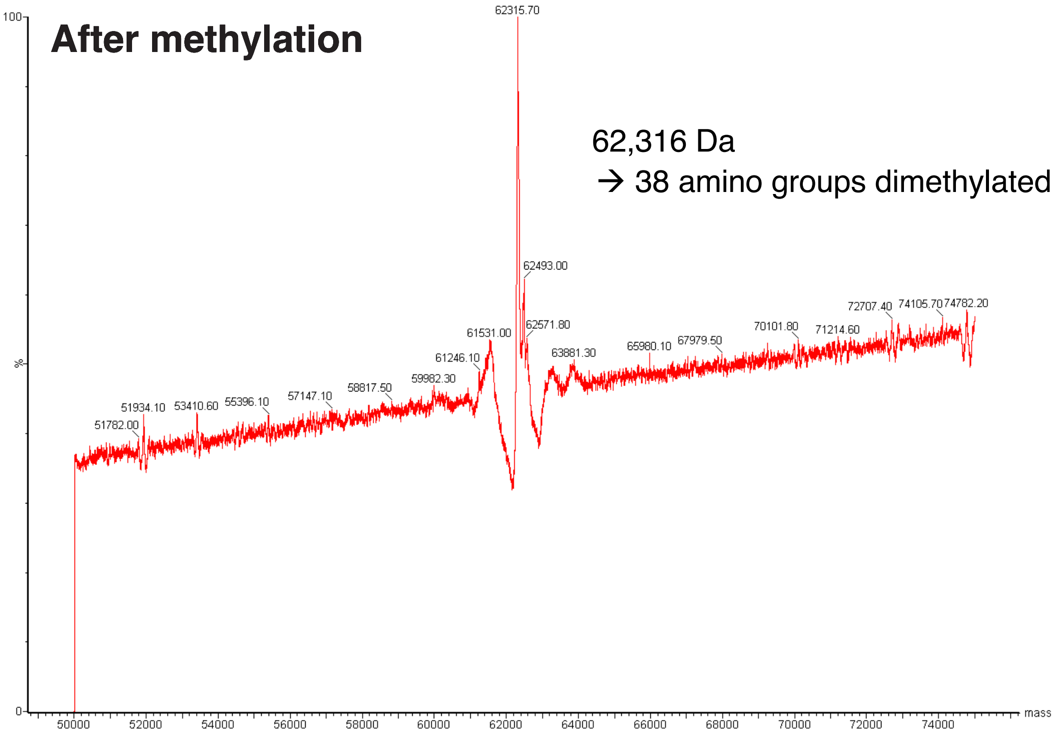


**Fig. S6.** Mass spectra of the T4L-KAP1 RBCC fragment before and after methylation of free amines (lysines and N-terminus) with dimethylamine borane complex (ABC).

**Table S1.** Crystallographic data collection and refinement statistics.

| **Data collection** | **T4 lysozyme-KAP1 RBCC** | | |
|---|---|---|---|
| X-ray source | DLS I03 | | |
| Space group | $C222_1$ | | |
| Cell dimensions | | | |
| a, b, c (Å) | 59.77, 169.3, 374.5 | | |
| α = β = γ (°) | 90 | | |
| Wavelength (Å) | 1.28189 | | |
| Resolution (Å) | 187–2.63 | 84.1–3.9[b] | 3.17–2.63 |
| Observations | 156,326 | 125,590 | 5,596 |
| Unique reflections | 33,980 | 15,929 | 924 |
| $R_{merge}$ [b] | 0.078 | 0.088 | 0.832 |
| $R_{pim}$ [c] | 0.029 | 0.034 | 0.363 |
| $\langle I \rangle / \sigma I$ | 13.4 | 8.14 | 2.2 |
| Spherical completeness (%) | 32.4 | 89.2 | 3.9 |
| Ellipsoidal completeness (%) | 86.6 | – | 61.7 |
| Multiplicity | 8.5 | 7.9 | 6.1 |
| CC(1/2) | 0.998 | 0.994 | 0.820 |
| **SAD Phasing** | | | |
| $CC_{ano}$ | 0.479 | | |
| $\lvert D_{ano} \rvert / \sigma D_{ano}$ | 0.918 | | |
| Overall figure of merit | 0.67 | | |
| **Refinement** | | | |
| Resolution (Å) | 63–2.9 | | |
| $R_{work}$ / $R_{free}$ [d] | 0.261 / 0.291 | | |
| No. of non-H atoms | | | |
| Protein | 6955 | | |
| $Zn^{2+}$ Ions | 8 | | |
| Solvent | 0 | | |
| No. riding H atoms | 6908 | | |
| Mean B-factor ($Å^2$)[e] | 111 | | |
| MolProbity Clashscore | 7.35 | | |
| RMS[f] deviations | | | |
| Bond lengths (Å) | 0.005 | | |
| Bond angles (°) | 0.859 | | |
| Ramachandran plot | | | |
| % favored | 93.2 | | |
| % allowed | 6.45 | | |
| % outliers | 0.35 | | |
| PDB code | PDB: 6QAJ | | |

[b]Dataset reprocessed at 84.1-3.9 Å resolution with CCP4 (MOSFLM, AIMLESS)

[b]$R_{sym} = \Sigma_{hkl}\Sigma_i |I_{hkl,i} - \langle I \rangle_{hkl}| / \Sigma_{hkl}\Sigma_i|I_{hkl,i}|$, where $I_{hkl}$ is the intensity of a reflection and $\langle I \rangle_{hkl}$ is the average of all observations of the reflection.

[c]$R_{pim} = \Sigma_{hkl} (N_{hkl} - 1)^{-1/2} \times \Sigma_i |I_{hkl,i} - \langle I \rangle_{hkl}| / \Sigma_{hkl}\Sigma_i|I_{hkl,i}|$, where $I_{hkl}$ is the intensity of a reflection and $\langle I \rangle_{hkl}$ is the average of all observations of the reflection.

[d]$R_{free}$, $R_{work}$ with 5% of $F_{obs}$ sequestered before refinement.

[e]Residual B-factors after TLS refinement. See PDB entry for TLS refinement parameters.

[f]R.M.S., root mean square.
